# Phospholipase D1 Produces Phosphatidic Acid at Sites of Secretory Vesicle Docking and Fusion

**DOI:** 10.1101/2023.12.07.570684

**Authors:** Broderick L. Bills, Megan L. Hulser, Michelle K. Knowles

## Abstract

Phospholipase D1 (PLD1) activity is essential for the stimulated exocytosis of secretory vesicles where it acts as a lipid-modifying enzyme to produces phosphatidic acid (PA). PLD1 localizes to the plasma membrane and secretory vesicles, and PLD1 inhibition or knockdowns reduce the rate of fusion. However, temporal data resolving when and where PLD1 and PA are required during exocytosis is lacking. In this work, PLD1 and production of PA are measured during the trafficking, docking, and fusion of secretory vesicles in PC12 cells. Using fluorescently-tagged PLD1 and a PA-binding protein, cells were imaged using TIRF microscopy to monitor the presence of PLD1 and the formation of PA throughout the stages of exocytosis. Single docking and fusion events were imaged to measure the recruitment of PLD1 and the formation of PA. PLD1 is present on mobile, docking, and fusing vesicles and also colocalizes with Syx1a clusters. Treatment of cells with PLD inhibitors significantly reduces fusion, but not PLD1 localization to secretory vesicles. Inhibitors also alter the formation of PA; when PLD1 is active, PA slowly accumulates on docked vesicles. During fusion, PA is reduced in cells treated with PLD1 inhibitors, indicating that PLD1 produces PA during exocytosis.

## INTRODUCTION

Regulated exocytosis is a tightly controlled process in neuroendocrine cells that is essential for the secretion of hormones and neurotransmitters. During exocytosis, the membrane of a docked vesicle fuses with the plasma membrane, a process with a high energy barrier. SNARE proteins have been shown to provide the minimal machinery for fusion (Weber *et al*., 1998) and a few copies, along with their accessory proteins, can provide the energy required (Mohrmann *et al*., 2010; Stepien and Rizo, 2021). While SNARE proteins are vital for exocytosis, lipid rearrangement has been proposed to assist by recruiting protein clusters (Lang *et al*., 2001), recruiting vesicles (Honigmann *et al*., 2013), or stabilizing the highly curved fusion pore (Chernomordik and Kozlov, 2005; Rohrbough and Broadie, 2005; McMahon *et al*., 2010). For example, phosphatidylinositol (4,5)-bisphosphate (PI (4,5)P_2_) has been shown to inhibit fusion pore dilation (Omar-Hmeadi *et al*., 2023) and cholesterol is involved with the clustering of Syntaxin1a, a plasma membrane associated SNARE protein essential to docking and fusion (Lang *et al*., 2001). Lipid-modifying enzymes, such as phospholipase D1 (PLD1), have been implicated as essential in exocytosis (Humeau *et al*., 2001; Vitale *et al*., 2001; Hughes *et al*., 2004; Jenkins and Frohman, 2005; Zeniou-Meyer *et al*., 2007) and its product, phosphatidic acid (PA), can stabilize negatively curved membranes (Kooijman *et al*., 2003; Chernomordik and Kozlov, 2005; Callan-Jones *et al*., 2011; Bills and Knowles, 2022).

There are six PLD isoforms in mammals, with PLD1 and PLD2 acting as lipases positioned within to play a role in exocytosis (Brown *et al*., 1998; Vitale *et al*., 2001; Cockcroft *et al*., 2002; Hughes *et al*., 2004; Jenkins and Frohman, 2005). PLD1 and PLD2 use the substrate phosphatidylcholine to produce PA (Frohman, 2015). PLD1 and PLD2 primarily differ in basal activity; PLD2 is constitutively active and PLD1 is stimulated (Peng and Frohman, 2012). Several studies have shown that inhibition and knockdowns of PLDs reduce exocytosis (Humeau *et al*., 2001; Vitale *et al*., 2001; Hughes *et al*., 2004; Jenkins and Frohman, 2005; Zeniou-Meyer *et al*., 2007). Specifically, PLD1 is essential for exocytosis in platelets, HL-60 cells, PC12 cells and chromaffin cells (Haslam and Coorssen, 1993; Stutchfield and Cockcroft, 1993; Zeniou-Meyer *et al*., 2007, 2009), where the loss or inhibition of PLD1 diminishes secretion, demonstrating that that PLD1 or PA is required. However, PLD2 has also been shown to be involved in exocytosis, particularly in mast cells, where PLD1 and PLD2 are involved at different stages (Choi *et al*., 2002). Overall, PLD1 is required for stimulated exocytosis in most secretory cells, and it is likely that the formation of PA is essential.

PLD1 localizes partially to the plasma membrane (Brown *et al*., 1998; Freyberg *et al*., 2001; Hozumi *et al*., 2022), and its activity increases during stimulation leading to the formation of PA at the plasma membrane (Zeniou-Meyer *et al*., 2007; Tanguy *et al*., 2020). PA formation has typically been observed by the recruitment of the PA binding domain (PABD) of Spo20 to the plasma membrane (Zeniou-Meyer *et al*., 2007, 2009; Tanguy *et al*., 2022). However, new dyes have been designed to show where PA is newly formed, using click chemistry and relying on PLD activity (Bumpus and Baskin, 2017). One hypothesis for the role of PA is during the membrane fusion step of secretion, where PA could stabilize the highly curved fusion pore (Kooijman *et al*., 2003; Chernomordik and Kozlov, 2005; Callan-Jones *et al*., 2011; Bills and Knowles, 2022). PA is an inverse conical lipid containing a small, negatively charged headgroup with two fatty acid tails. PA has been established as preferring negative curvature in *in vitro* studies (Kooijman *et al*., 2003, 2005; Bills and Knowles, 2022). However, precisely when and where PA is formed during exocytosis is currently lacking.

To test if PLD1 is present at fusing vesicles and determine when and where the production of PA is required, a model secretory cell line (PC12) was used to express either GFP-PLD1 or GFP-PASS, a PA biosensor similar to PABD (Zhang *et al*., 2014). Cells were imaged using total internal reflection fluorescence microscopy and the presence of PLD1 and the formation of PA were compared with the location of secretory proteins: Syntaxin-1a (Syx1), vesicle associated membrane protein 2 (VAMP2) and neuropeptide Y (NPY). Single fusion events were imaged at high speed to measure the recruitment of PLD1 and the formation of PA as vesicles docked and fused with the plasma membrane. Colocalization analyses under basal, stimulated, and inhibitory conditions demonstrate that PLD1 is present on docked, fusing, and moving vesicles and colocalizes with Syx1 clusters. The presence of PLD1, however, does not solely determine where PA is produced. Although PLD1 is present on vesicles, the production of PA only occurs after vesicles dock and after membrane fusion. The production of PA at these stages requires PLD1.

## RESULTS

### PLD1 localizes to secretory vesicles and Syx1 clusters

To determine the role of PLD1 in SV secretion, PC12 cells were transiently transfected with GFP-PLD1 and either VAMP2-pHmScarlet, NPY-mCherry or Syx1-mCherry. GFP-PLD1 visually colocalizes to all three proteins (Figures. 1A-C). To measure the extent of colocalization, an object-based analysis was used where locations of interest, such as Syx1a clusters and secretory vesicles, were located and the GFP-PLD1 intensity was measured at these sites. To quantify the amount of GFP-PLD1 at sites of interest, the intensity within a circle centered around the vesicle or Syx1 cluster location was measured relative to the local background (ΔF = circle – annulus, shown in Figure 1D). This was normalized by the expression level (S = annulus – background outside of the cell). A positive value indicates that GFP-PLD1 is present at VAMP2 containing vesicles, NPY vesicles, or Syx1 clusters. GFP-PLD1 significantly localizes to all three compared to a negative control: cytosolic GFP. Upon stimulation with high K^+^ buffer, Ca^2+^ enters PC12 cells and fluorescence from a calcium indicator increases (Figure S1). All cells were labeled with CellMask (Figure S1A) and Fluo4-AM (Figures S1B-C). The only cells that did not show a change indicating the influx of Ca^2+^ were cells that were part of a larger cluster of cells (Figures S1D-E), therefore, single cells or pairs of cells were measured for the remaining experiments. No increase in colocalization was observed for PLD1 at VAMP2 positions after 2 minutes of stimulation (Figure 1D). To further confirm PLD1 localization, the JACoP plugin in ImageJ was used to identify the percentage of PLD1+ vesicles and Syx clusters, which are all significantly higher than cytosolic GFP at VAMP2 vesicles (Figure 1E). Additionally, to verify that green bleed through into the red channel was not causing the observed colocalization, zoomed in locations of cells expressing GFP-PLD1 and a red marker were examined. Many green spots do not show a red spot in the same position (Figure S1F). Overall, PLD1 is positioned to play a role in is exocytosis, in line with what others have shown (Brown *et al*., 1998; Vitale *et al*., 2001; Cockcroft *et al*., 2002; Hughes *et al*., 2004), and stimulation does not alter the position of PLD1 over a short period of time.

**Figure 1:**
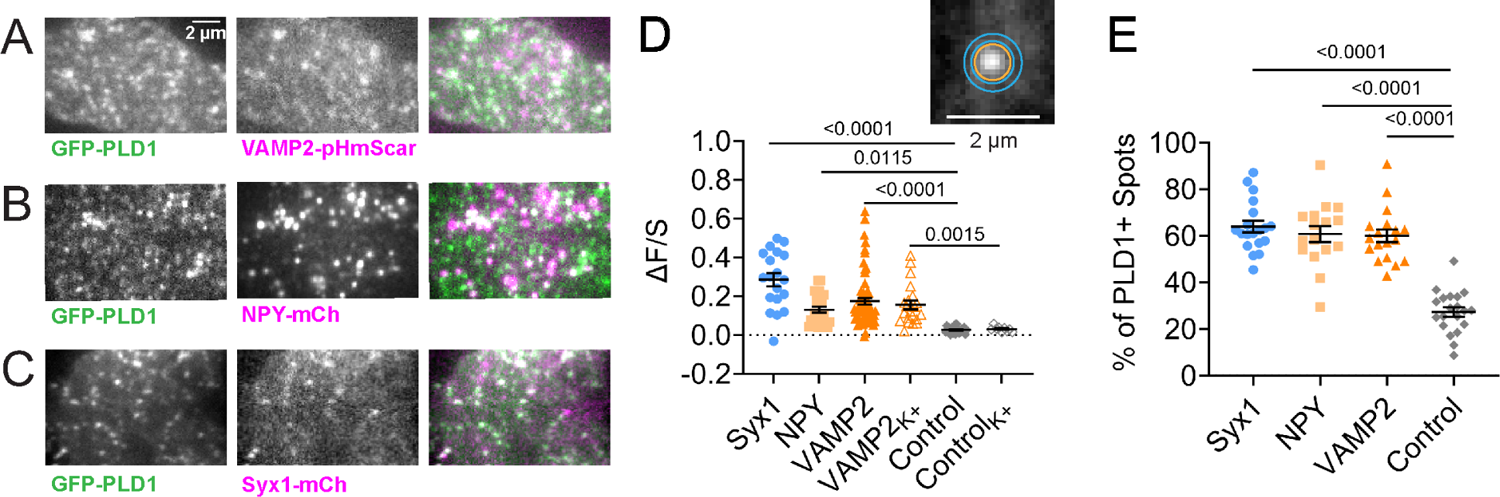
Phospholipase D1 Colocalizes with Secretory Vesicle Markers and Syntaxin Clusters. A-C) PC12 cells were transfected with GFP-PLD1 (left) and A) VAMP2-pHmScarlet, B) NPY-mCherry or C) Syx1-mCherry (middle) and imaged at room temperature using TIRF microscopy. The overlays (right) are white where colocalization occurs. Scale bar: 2 µm. D) The extent of GFP-PLD1 accumulation, ΔF/S, at Syx1 clusters (blue circles), NPY spots (light orange squares) or VAMP2 spots (dark orange triangles). As a control, ΔF/S was measured for cytosolic GFP at VAMP2 spots (grey diamonds). Empty symbols represent ΔF/S of PLD1 or cytosolic GFP in cells stimulated with 60 mM KCl. Lines and errors are mean and SEM. Black bars and numbers above represent p-values between PLD1 at sites of the indicated protein compared to cytosolic GFP at VAMP2+ vesicles. Each spot represents one cell. At least 10 cells from at least 3 independent experiments were conducted for each condition. *Inset*: description of ΔF/S measurements, where 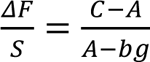. The circle, C, is represented by the orange circle, while annulus A is the space between the cyan circles and bg is the average intensity surrounding the cell. E) The JACoP plugin in ImageJ was used to identify the percentage of Syx1a or vesicle locations with PLD1 or cytosolic GFP. All statistics are described in the Methods.

One limitation of using GFP-PLD1 is that some studies suggest that overexpression of PLD1 leads to mislocalization (Freyberg *et al*., 2001). By eye, GFP-PLD1 is extremely low expressing as noted by how dim cells were on the microscope, suggesting that transient expression in PC12 cells is low. To quantify the amount of overexpression a slot blot of PLD1 from transiently transfected cells was compared to endogenous PLD1 expression (Figure S2A). PLD1 is increased by 4.3% in a Western blot of all cells (transfected and non-transfected), where about 25% of cells are transfected (Figure S2), therefore it is overexpressed by approximately 17% in transfected cells. Additionally, a different PLD1 construct with an internal GFP label has been created with the goal of moving the GFP moiety away from membrane binding motifs (Corrotte *et al*., 2006.). To determine if the colocalization that was observed with GFP-PLD1 (Figure 1) was also present with endogenous PLD1, immunofluorescence was performed on fixed cells; anti-PLD1 is present at sites of VAMP2-pHluorin (Figure S2D). This suggests that the N-terminal GFP tag and overexpression is not altering the position of PLD1 in PC12 cells.

### GFP-PLD1 is present on mobile VAMP2 vesicles and during docking, fusion

To assess whether PLD1 is recruited as vesicles traffic to the plasma membrane, dock and then fuse, cells expressing GFP-PLD1 and VAMP2-pHmScarlet were imaged in time using TIRF microscopy. VAMP2-pHmScarlet is visible prior to fusion and also increases in fluorescence due to the large pH change that occurs upon fusion (Liu *et al*., 2021), making it an excellent probe for visualizing moving, docking, and fusing vesicles in a single-color channel. VAMP2-pHmScarlet vesicles were located using a previously established method (Mahmood *et al*., 2023), then divided into three classes: visiting, docking, or fusing vesicles based on whether they move through a cropped movie (Figure 2A), appear and remain static (Figure 2B), or appear, increase quickly in intensity and spread out from the center (Figure 2C). To quantify the data, the intensity within a circular region (Figure 1D, inset orange) was measured in time and the average of five frames prior to the event onset (−0.5 to −0.1 s) was subtracted and normalized (see Methods). Note that the annulus was not used in this measurement because it changes in intensity when fusion occurs as fluorescence radially expands. The whole trace was then normalized by the maximum intensity. To visualize this change, events were averaged into one movie and then 5-frames at fusion and visitor onset (−0.5 to −0.1 s, “initial”) and VAMP2 peaks or plateaus (“final”) were averaged (Figures 2D-F, top row). To highlight the change that occurs in the GFP-PLD1 intensity during the events, the initial image was subtracted from the final image to create a difference image (“Δ”). All GFP-PLD1 difference images show spots in the center. As a control, cytosolic GFP was expressed in place of GFP-PLD1 (Figures 2D-F, bottom row). The intensity in time is shown for both the vesicle (Figures 2G-I, purple) and PLD1 (Figures 2G-I, green) such that 0 s is the beginning of the rise in VAMP2 intensity and the onset of the vesicle fusion, docking or visiting. In all three types of events, the intensity of GFP-PLD1 significantly increases after the onset of visiting, docking or fusion (Figures 2D-I), suggesting that PLD1 is carried on VAMP2 vesicles. Interestingly, during fusion (Fig 2I), the loss of PLD1 occurs faster than the loss of VAMP2 from the fusion site, possibly due to different rates of diffusion on the membrane. In all types of events, cytosolic GFP was used as a control and an increase in intensity is not observed (Figures 2G-I, S3). This suggests that PLD1 is present on moving VAMP2 vesicles and more may be recruited during docking and fusion.

**Figure 2:**
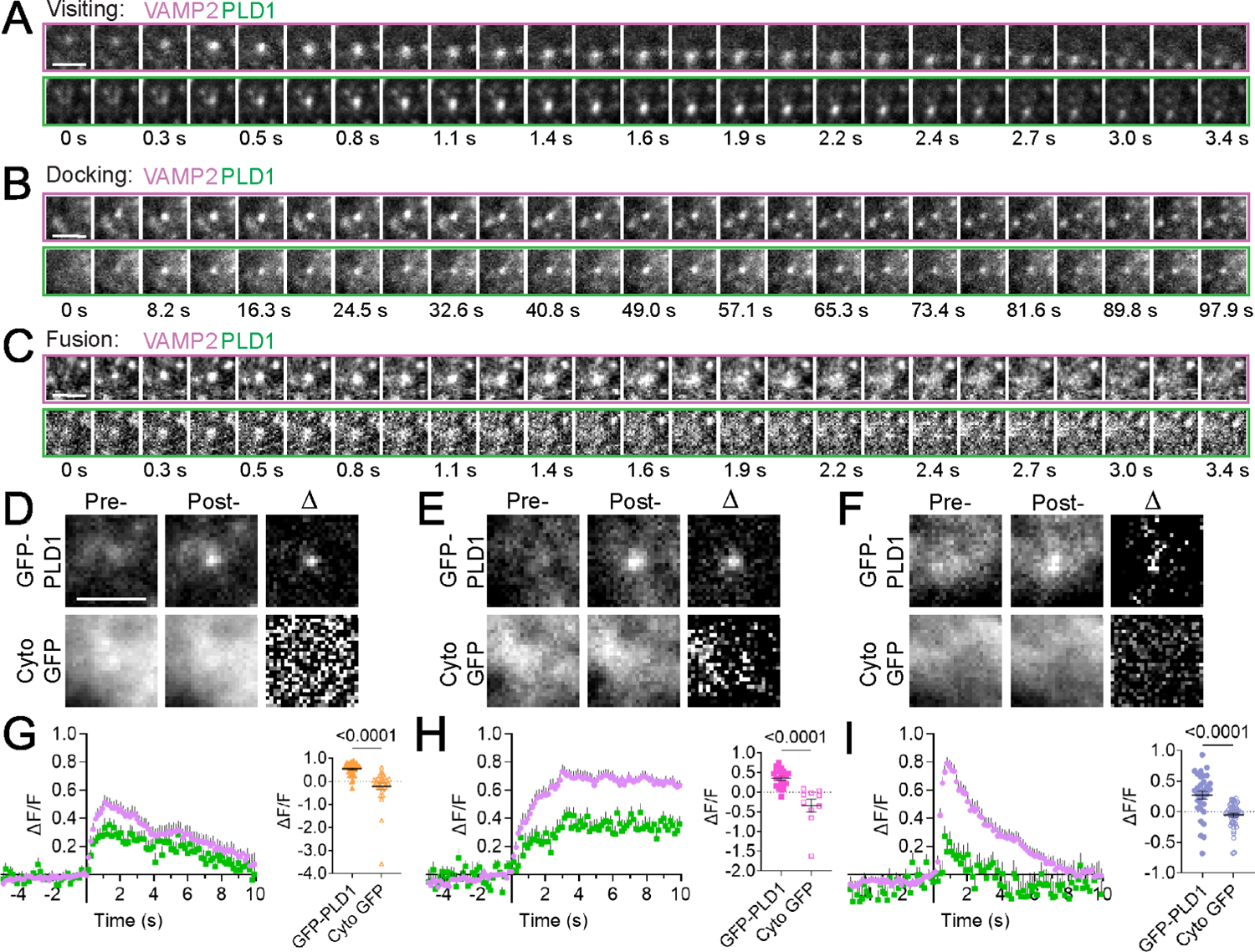
GFP-PLD1 Localizes to Visiting, Docking, and Fusing VAMP2 Vesicles. PC12 cells expressing GFP-PLD1 and VAMP2-pHmScarlet were imaged at 136 ms/frame at room temperature. A-C: Single vesicle events were located using an automated algorithm that identifies three types of events: visiting (A), docking (B) or fusing (C) vesicles Top rows: montages of VAMP2-pHmScarlet. Bottom rows: corresponding montages of GFP-PLD1, where each image is a 5-frame average. Scale bars: 2 µm. A-C are examples of a single event. D-F) Average images of many events prior to visiting (D, 17 PLD1 and 22 GFP events), docking (E, 18 and 10 events) or fusion (F, 22 and 25 events) at the onset (left) and peak/plateau VAMP2 intensity (middle) for PLD1 (top) or cytosolic GFP (bottom). A difference image, Δ, of each is shown to highlight changes (right). Initial and final images are contrasted the same. Scale bar: 2 µm. G-I) Left: Traces of G) visiting (n = 37), H) docking (n = 20) and I) fusing (n = 32) vesicles for GFP-PLD1 (green) and VAMP2-pHmScarlet (purple). The intensity, ΔF/F, was normalized to 1.0 for each event prior to averaging. Graphs depicting ΔF/F values of GFP-PLD1 (solid symbols) and cytosolic GFP (empty symbols) at the peak of the VAMP2 traces or at 80% of the plateau. Each spot represents one event for cytosolic GFP (7 cells, 3 replicate experiments) or GFP-PLD1 (17 cells, 6 replicate experiments). Line and error are mean and SEM; all statistics are described in the Methods.

### Inhibition of PLD1 reduces the fusion of VAMP2 vesicles and vesicle mobility but does not alter PLD1 localization

One key feature of PLD1 is its ability to convert PC into PA, where the lipid PA is then involved trafficking (Brito de Souza *et al*., 2014; Luo *et al*., 2017; Tanguy *et al*., 2022) and fusion (Humeau *et al*., 2001; Vitale *et al*., 2001; Hughes *et al*., 2004; Jenkins and Frohman, 2005; Zeniou-Meyer *et al*., 2007). To explore the activity of PLD1, PC12 cells expressing GFP-PLD1 and VAMP2-pHmScarlet were transfected with control or PLD1 siRNA. Fusion rates were reduced by ∼70% under knockdown conditions, which was not rescued by transfecting cells with GFP-PLD1 (Figure 3A). This rescue possibly failed due to the very low expression of GFP-PLD1 in PC12 cells (Fig S2). Further testing was conducted with PLD inhibitors, either 100 nM FIPI, a pan-PLD inhibitor or 500 nM VU0155069, a PLD1 specific inhibitor. To verify if fusion is blocked in the presence of inhibitors, the rate of fusion was measured. The frequency of fusion was significantly reduced in K^+^ stimulated cells when cells were treated with siRNA (Figure 3A). PLD1 and PLD1/2 inhibitors (Figure 3B) reduced fusion to a similar extent, however, the localization of GFP-PLD1 to sites of fusion was not significantly affected (Figure 3C). The inhibition of PLDs also altered the rate vesicles near the plasma membrane moved. VAMP2 vesicles were tracked in time and the rate of motion was measured. Diffusion coefficients of VAMP2+ vesicles in cells treated with inhibitors were slightly slower (Figure S4A). However, when only mobile vesicles (D > 0.0055 µm^2^/s) were counted (Figure S4B), the reduction in motion vanishes, suggesting that the reduction observed is due to a change in the number of mobile vesicles. Interestingly, the fraction of mobile vesicles per cell does not significantly decrease upon treatment with either inhibitor (Figure S4C). Therefore, the activity of PLD1 impacts the mobility and fusion of VAMP+ vesicles but does not contribute to the location of PLD1.

**Figure 3:**
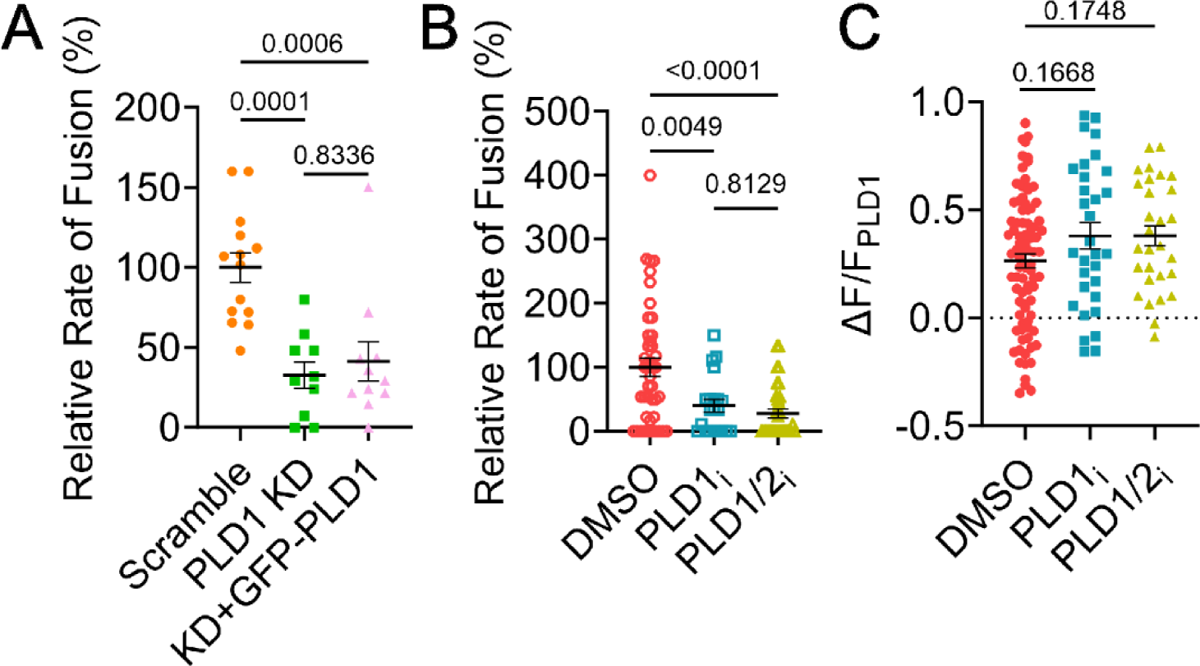
PLD Knockdown and Inhibition Reduces Fusion Rates. A) PC12 cells were treated with siRNA (Scramble, orange circles or PLD1 siRNA, green squares and pink triangles) and expressed VAMP2-pHmScarlet (orange circles and green squares) or VAMP2-pHmScarlet and GFP-PLD1 (pink triangles). Fusion events were identified and counted in cells stimulated with 60 mM KCl, then divided by the average rate of fusion in Scramble cells to account for day-to-day variation. Each point represents one cell from 3 independent experiments. B) PC12 cells expressing VAMP2-pHmScarlet and GFP-PLD1 were treated with inhibitors for PLD1 (VU0155069, 500 nM), PLD1 and 2 (FIPI, 100 nM), or a similar amount of DMSO (control). Cells were incubated 30 min prior to imaging and imaged at 37°C. The relative frequency of VAMP2 fusion events decreases with inhibition. This includes spontaneous and evoked fusion. Each point is one cell from at least 4 independent experiments. C) The intensity (ΔF/F) of PLD1 at peak VAMP2 intensity during fusion in DMSO, PLD1i or PLD1/2i treated cells. Each point is one fusion event. Lines are mean values, error bars are SEM. All statistics are described in the Methods.

### PA accumulates at fusion sites and this depends on PLD1 activity

If PLD1 activity is essential to fusion, the formation of PA should occur and the timing of PA formation during the fusion process can be determined by imaging single fusion events. To test this, fusion events were located in cells expressing VAMP2-pHmScarlet and GFP-PASS analogous to the approach shown for GFP-PLD1 (Figure 2). GFP-PASS is a PA binding protein tagged to mGFP and marks regions of the cell that are enriched in PA (Zhang *et al*., 2014). Visiting, docking and fusion events were identified, and average GFP-PASS images were calculated from events prior to visiting (Figure 4A), docking (Figure 4B) or fusing (Figure 4C) vesicles. Figure 4 shows the PASS accumulation only and PASS does not show up during the initial 2-3 seconds over which the images are shown for vesicles that move (Figure 4A) or dock (Figure 4B). However, during fusion, GFP-PASS increases quickly (Figure 4C). A difference image visualizes the change in PA via the GFP-PASS intensity in all three classes of events (Figures 4A-C). To quantify the intensity change, the ΔF/F of PASS during visiting, docking and fusion were calculated (Figures 4D-F). Unlike PLD1, PASS does not significantly increase immediately upon visiting or docking but does increase during fusion (Figure 4G), reaching a maximum 1.4s after the peak of the VAMP2 fusion event (Figure 4H).

**Figure 4:**
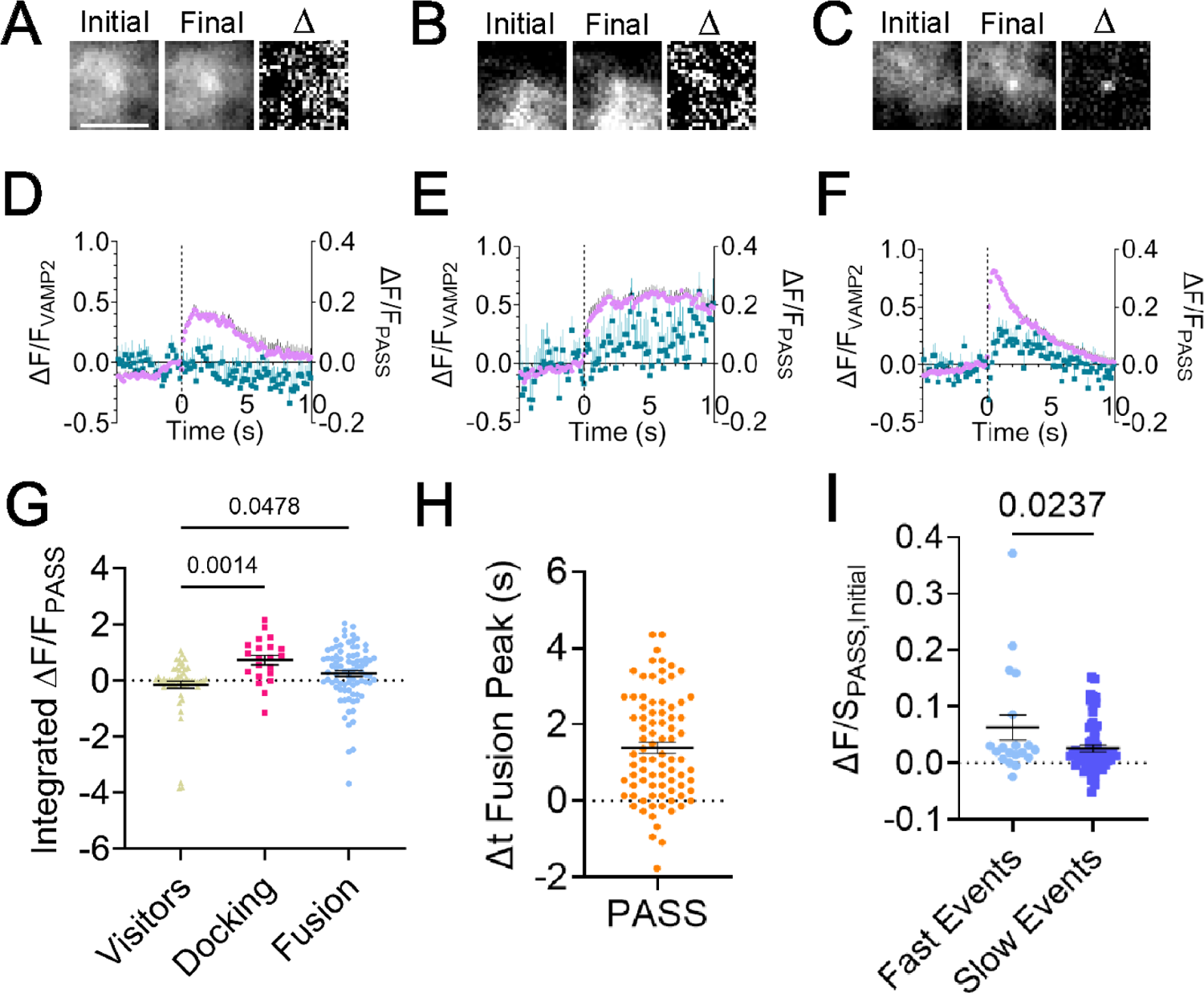
PA Is Present During Fusion and Docking. PC12 cells were transfected with GFP-PASS, a PA binding protein, and VAMP2-pHmScarlet, then cells were imaged to visualize A) visiting, B) docking and C) fusing vesicles. A-C) Average images of GFP-PASS prior to onset (left), at the peak (middle) of the VAMP2 intensity, and a difference image between the two (right). Scale bar: 2 µm. Before and after images are contrasted identically and are averages of 5 frames (n=51, 12 and 46 for A-C, respectively). D-F) Average intensity (ΔF/F) traces of VAMP2-pHmScarlet (purple circles) and GFP-PASS (teal squares). Error bars are SEM (n = 62, 21, 83 events for D-F, respectively, from at least 8 cells and 3 independent experiments). G) Integrated intensity of GFP-PASS from 0 to 4 seconds after the fluorescence onset of VAMP2 during visiting, docking or fusing events. Each point represents one event from 3 independent experiments. H) Time from peak VAMP2 intensity to peak PASS intensity. Each point represents one event. I) Initial VAMP2 decay slopes were binned into fast and slow events and the amount of PASS present was measured for each. Fast events have more PASS present. G-I) Lines are mean values, error bars are SEM; all statistics are described in the Methods.

PA has been hypothesized to accumulate at negatively curved regions within the fusion pore, which suggests that the amount of PA could possibly affect the release rate of content post-fusion. To probe whether the presence of PA affects the fusion kinetics, the slope of the decay of VAMP2 from the peak fluorescence to 1 s later and the GFP-PASS intensity (ΔF/S) was measured for single fusion events. It is useful to note that the measurement ΔF/S is normalized by the local background (S) and conveys enrichment at the vesicle position, and this is insensitive to the expression level within a range (Barg *et al*., 2010). The initial ΔF/S of GFP-PASS was plotted against the initial decay of the VAMP2 intensity post-fusion (Figure S5) and a negative correlation was noted (Pearson’s correlation coefficient = −0.1744), which indicates a higher PA presence during faster fusion events. Therefore, the rate of decay was binned into “fast” and “slow” events based on the histogram of slopes (Figure S5). There is more GFP-PASS present on faster events (Figure 4I).

To probe whether PA is produced by PLD1 during fusion rather than recruited or produced by other means, PC12 cells expressing GFP-PASS and VAMP2-pHmScarlet were treated with inhibitors for PLD1 or PLD1/2 and compared to a vehicle control, as previously described (Figure 3). Cells were stimulated with K+ during imaging. Due to the fact that PLD inhibitors reduce fusion events (Figures 3A-B), the number of events observed in cells treated with inhibitors was low. GFP-PASS intensity from fusion locations were averaged before and during fusion, and the difference between the two was measured (Figure 5A-C, *inset*). The average fusion traces were plotted (Figure 5A-C, purple) alongside the GFP-PASS intensity (Figure 5A-C, blue). To quantify whether PLD1 inhibition altered the recruitment of GFP-PASS during fusion, the change in GFP-PASS intensity was measured at the time that coincided with the peak of the VAMP2-pHmScarlet intensity relative to the pre-fusion intensity (Figure 5D). All inhibitory conditions show a decrease in PASS intensity, possibly due to new membrane lacking GFP-PASS being added to the fusion site. However, GFP-PASS increases in intensity in control conditions and this is significantly more than when PLD is inhibited (Figure 5D). Because these inhibitors do not affect other PA-producing enzymes, like diacylglycerol kinase, this reduction of GFP-PASS intensity is likely due to PLD1 and the lack of PA.

**Figure 5:**
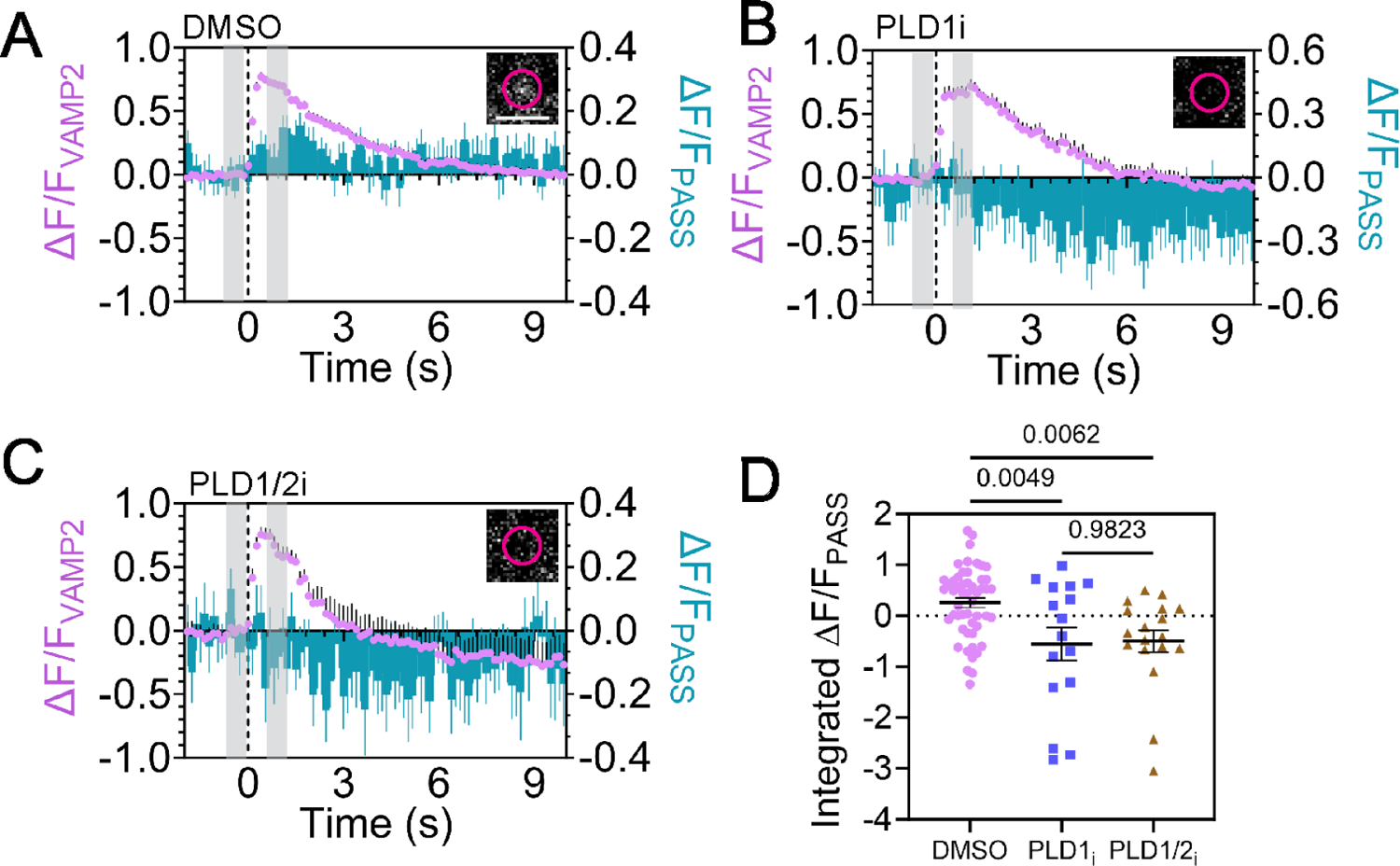
Inhibition of PLD1 reduces the amount of PA present during fusion. PC12 cells were transfected with GFP-PASS (a PA marker) and VAMP2-pHmScarlet, then cells were imaged at 37°C to visualize fusing vesicles in the presence of PLD inhibitors. A) Control cells treated with DMSO (n = 40 cells, 52 fusion events), B) Cells treated with PLD1 specific inhibitor (n = 12, 16 fusion events), and C) Cells treated with a pan-PLD inhibitor (n = 13, 19 fusion events). *Inset:* Average image showing the change in PASS intensity, where the average image from five frames immediately prior to fusion was subtracted from the average PASS image (5 frames) that coincides with the VAMP2 peak. The time regions are shown in the grey rectangles and the scale bar is 2 µm. In A-C the average normalized intensity traces of fusion events is shown for VAMP2-pHmScarlet (purple) and GFP-PASS (cyan). D) Integrated PASS intensity from 0 to 4s after the onset of the VAMP2-pHmScarlet fusion event. Lines are mean values, error bars are SEM; all statistics are described in the Methods.

### PA accumulates at vesicle locations after docking and throughout exocytosis

PA is present near vesicle docking sites in EM data (Zeniou-Meyer *et al*., 2007) and accumulates on the plasma membrane minutes after stimulation (Tanguy *et al*., 2020). To determine the time course of PA arrival after vesicles dock, cells were stimulated with high K^+^ and GFP-PASS was imaged and quantified during docking for 10s of seconds after the initial docking event (Figures 6A-B). Difference images show the increase of PASS at the docking cite (Figure 6A, bottom). After 10s, the GFP-PASS signal is significantly larger than the cytosolic-GFP control (Figure 6C), and the GFP-PASS signal gradually increases as time goes on (Figure 6B). To identify if PA is formed or accumulated during vesicle docking, PLD inhibitors were used and both inhibitors block the formation of PA (Figure 6D), therefore PA is formed by PLD1 during docking events and PLD2 cannot compensate for PLD1 inhibition. When the intensity of GFP-PASS is measured relative to the amount present prior to docking (considered a non-specific, background fluorescence because GFP-PASS is present in the cytoplasm), the intensity of GFP-PASS increases slowly after docking, then more prior to fusion, reaching a maximum post-fusion (Figure 6E). If cytosolic GFP is expressed in place of GFP-PASS, this intensity increase is not observed (Figure S6). Therefore, PA formation increases throughout the entire exocytosis process, from moving to docking to fusion, as exocytosis progresses. This places PA in a position to regulate several stages of membrane fusion.

**Figure 6:**
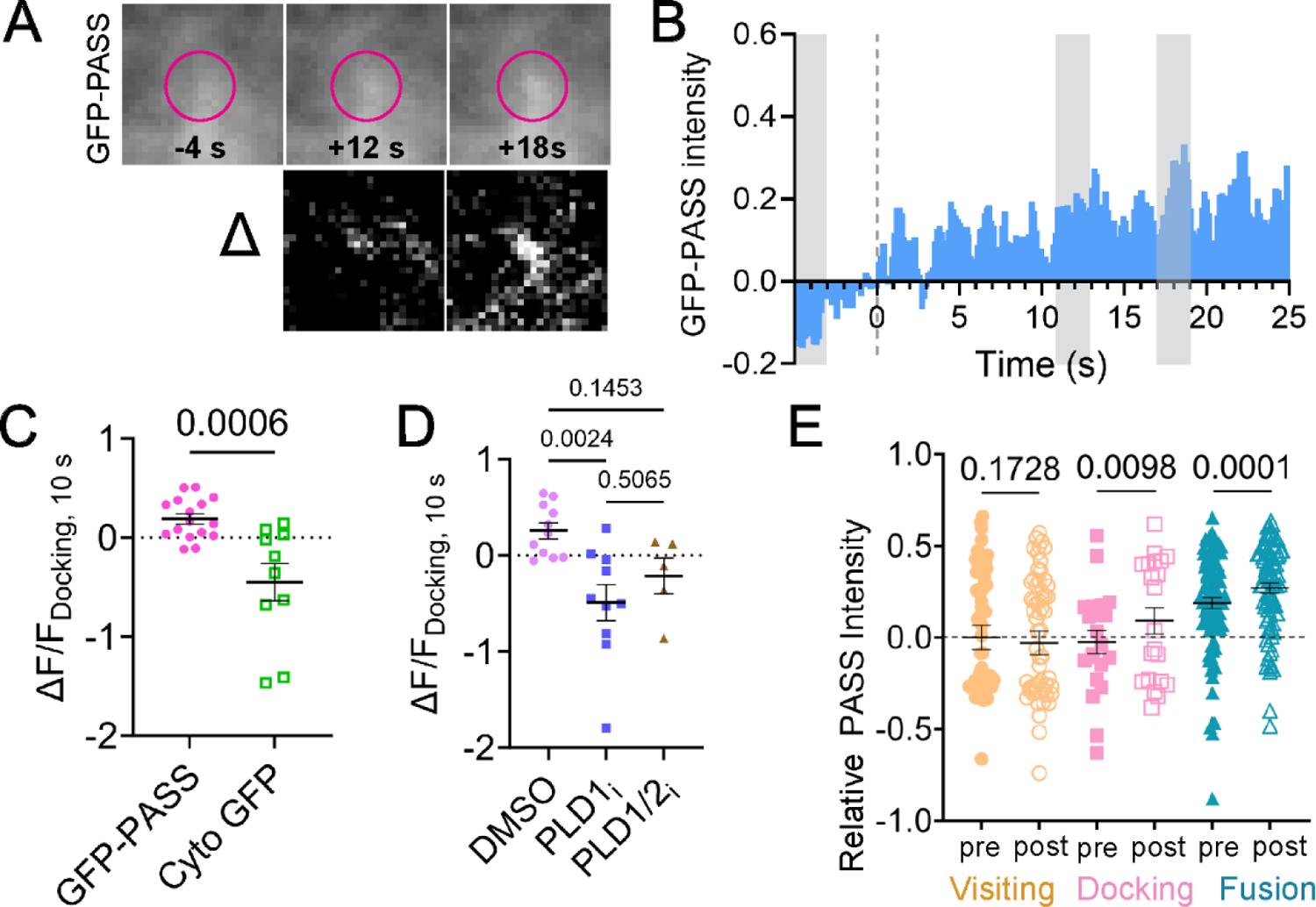
PLD1 Produces PA at Docked Vesicles over Time in Stimulated Cells. A) Average images (n = 13 events) of PASS before onset of docking (−4s), after 8 s, and after 12 s post-docking (top). Pink circles mark the location of the docked vesicle. The difference between pre and post docking images (bottom). Initial and final images are contrasted identically. B) The average ΔF/F trace of GFP-PASS over extended periods of time after stimulation with 60 mM KCl. The grey boxes refer to the times shown in A. C) ΔF/F of PASS (pink circles) and GFP (green squares) 10 s after docking. One dot corresponds to one docking event. D) ΔF/F of PASS 10 seconds after docking in cells treated with DMSO (purple circles) with or without PLD1 inhibition (blue squares) or PLD1/2 inhibition (brown triangles). E) The relative PASS intensity was measured over the different stages of membrane fusion: visiting (orange), docking (pink), and fusion (blue). The PASS pre intensity (visiting) was subtracted from all to obtain a relative intensity. The “pre” intensity was measured at 0.5 to 0.1 s prior to the event. The post-docking intensity (pink, open circles) was measured after several seconds, when single events reached the plateau intensity in the VAMP2-pHmScarlet channel. The post-fusion intensity (blue, open circles) was measured when the VAMP2-pHmScarlet intensity was at a maximum. Lines in C-E are mean values, error bars are SEM. All statistics are described in the Methods.

## DISCUSSION

In this work, we demonstrate that the production of PA via PLD1 activity impacts exocytosis in PC12 cells. The inhibition of PLD1 or PLD1 and 2 reduces fusion events (Figures 3A-B). Specifically, PLD1 is required for fusion and the presence of PLD2 cannot compensate for the inhibition of PLD1 in PC12 cells (Figure 3). This agrees with the work of others, where the loss of PLD1 function by inhibitors, siRNA, or KO reduces stimulated secretion (Tanguy *et al*., 2022) and the frequency of fusion events in chromaffin cells (Tanguy *et al*., 2020). A wide variety of cells, such as adipocytes (Huang *et al*., 2005), endothelial cells (Disse *et al*., 2009), mast cells (Choi *et al*., 2002) and neuroendocrine cells (Zeniou-Meyer *et al*., 2007, 2009), need PLD1 for proper secretion, albeit some cell types (mast cells) also use PLD2 during different stages of exocytosis (Choi *et al*., 2002).

By imaging single secretory vesicles concurrent with PLD1, PLD1 can be shown to localize with secretory vesicles, as well as Syx1 clusters under basal conditions, suggesting an association between PLD1 and secretory machinery (Figure 1). This places PLD1 in a position to act when needed. As single vesicles were observed to visit the plasma membrane or dock, PLD1 intensity significantly increased (Figures 2A-B, D-E, G-H), indicating that PLD1 is trafficked on secretory vesicles. In regard to docking and visiting vesicles, the PLD1 intensity increase could be attributed to an increase in excitation as the vesicle moves into the TIRF field or an increase in the amount of PLD1, and these conclusions are challenging to disentangle. Therefore, we conclude that PLD1 is on vesicles as they dock and move near the plasma membrane and these results align with the established role of PLD1 in trafficking (Brito de Souza *et al*., 2014; Luo *et al*., 2017; Tanguy *et al*., 2022) and exocytosis (Humeau *et al*., 2001; Vitale *et al*., 2001; Hughes *et al*., 2004; Jenkins and Frohman, 2005; Zeniou-Meyer *et al*., 2007).

One main function of PLD1 is to catalyze the production of PA from PC, where the formation of PA has been shown to interact with the polybasic region of Syx1a clusters (Lam *et al*., 2008) and hypothesized to stabilize negative curvature within the fusion pore (Zeniou-Meyer *et al*., 2007; Gasman and Vitale, 2017). The role of PA and positioning of PLD1 is depicted in Figure 7 for docking, fusing, and visiting vesicles. Through high spatial and temporal imaging of single vesicles, the formation of PA was observed after vesicles stably dock (Figures 6A-C) and this formation requires PLD1 (Figure 6D). PASS is not present on vesicles that merely visit the plasma membrane (Figures 4A-D) and the GFP-PASS intensity does not immediately increase as vesicles dock (Figures 4B-E), suggesting that PA is not carried on vesicles. However, the rate that PASS binds to PA is a factor that needs to further examined to better understand the timing of PA formation. Others have noted that PA is enriched on vesicles (Kassas *et al*., 2017). It is possible that PA is on the inner leaflet or PASS is restricted from vesicular PA, like others have noted with the related probe, PABD of Spo20 (Carmon *et al*., 2020). Vesicles dock and PA slowly accumulates (Figures 6A-C). It is not clear, in our work, whether PA formation occurs on the vesicle membrane or the plasma membrane, but ultrastructural studies show the accumulation of PA near or on the plasma membrane near docked vesicles (Zeniou-Meyer *et al*., 2007). PA accumulation at the plasma membrane post stimulation has also been observed in PC12 cells using cellular fractionation and mass spectrometry methods (Zeniou-Meyer *et al*., 2007; Tanguy *et al*., 2020) and an increase in PA at the plasma membrane was also observed using fluorescence microscopy (Zeniou-Meyer *et al*., 2007). Together, this suggests that PA forms on the plasma membrane after docking. Similar to other fusion regulatory molecules, PA is not present in a cluster prior to docking; Syx1a clusters form after vesicles approach the membrane and Syx1a clusters are required for stable docking (Barg *et al*., 2010; Knowles *et al*., 2010). PA also accumulates after vesicles dock, and we hypothesize that PA could be retained at the docking site via the established interaction with Syx1a (Lam *et al*., 2008).

**Figure 7:**
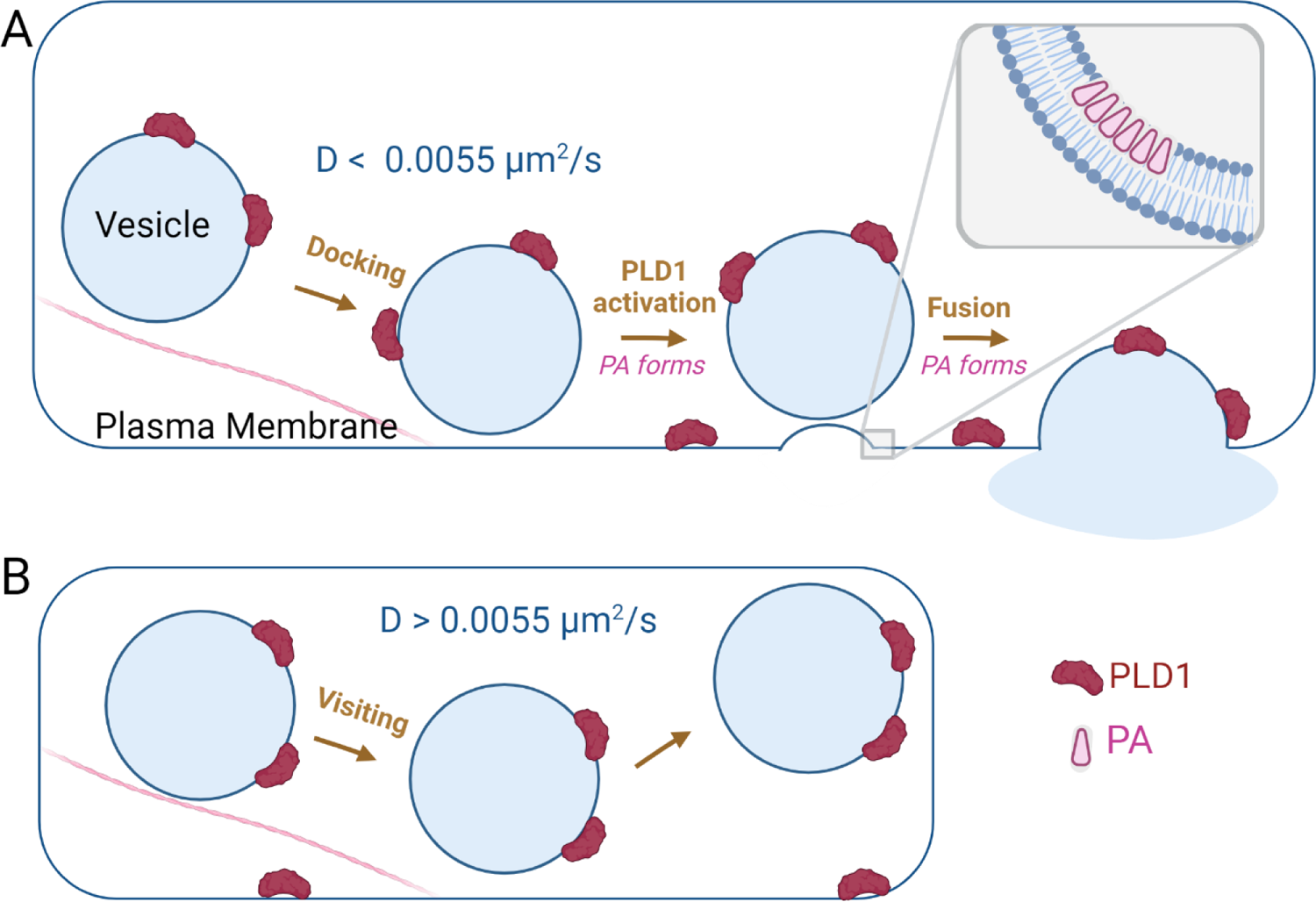
Model of PLD1 activity and localization. A) During docking and fusion, PA forms post-docking over the course of 10s of seconds and post-fusion, within seconds. *Inset:* A zoomed in look at the hypothesized role of PA pre-fusion at the grey box shown. PA has been shown to induce or sort at regions of negative membrane curvature and is hypothesized to assist with the membrane shape changes during fusion. B) Visiting secretory vesicles carry PLD1 but PA is not produced during movement. Moving vesicles move at a diffusion rate of 0.0055 µm^2^/s or higher.

We initially expected an increase in immobile/docked vesicles upon PLD KD or inhibition because fusion was inhibited (Figures 3A-B), therefore vesicles should be waiting at the membrane, unable to secrete. However, the number of immobile vesicles was not significantly different with inhibition (Figure S4A). This supports the idea that PA production at the docking sites aids in stabilizing the protein machinery necessary for docking. This could occur through proteins that are known aid in docking and interact with PA, such as Syx1a. Syx1a clusters are stabilized by PA (Lam *et al*., 2008), essential for docking in neuroendocrine cells (Knowles *et al*., 2010), and PLD1 accumulates at Syx1a clusters (Figure 1D). Others have noted that a longer treatment of high K^+^ during PLD inhibition reduced docked vesicles in EM data (Tanguy *et al*., 2020), further supporting the model that PA forms due to PLD activity at vesicle docking sites. Overall, our findings suggest PA formation could be a hallmark of docked vesicles, possibly acting through Syx1a.

After secretory vesicles initially dock, the amount of PA at docked vesicle positions gradually increases (Figure 6B) and continues to increase throughout the stages of exocytosis where PA is present immediately prior to fusion (Figure 6E). PA then increases post-fusion (Figures 4C, 4F, 6E) where GFP-PASS intensity hits a maximum at 1.4 s post fusion, on average (Figure 4H). To determine the source of PA, PLD1 and PLD1 and 2 were inhibited (Figure 5). Both inhibitors stop the accumulation of PASS at fusion sites, demonstrating that PLD1 is responsible for PA that accumulates post-fusion and the presence of PLD2 cannot compensate. PA could form *in situ* at the vesicle location or be recruited from PLD1 formed PA that is already present on the plasma membrane. Both potential sources of PA require PLD1 and cannot be compensated with PLD2 (Figure 5).

It is interesting to note that PLD1 is present in places where PA is not observed. For example, PLD1 is on both moving and docking vesicles (Figure 2), yet PA forms post-docking and post-fusion. This suggests that PLD1 is waiting for activation to begin PA production. PLD is activated by the V-ATP synthase subunit V0a1 and this interaction requires ARNO, a GEF protein for Arf6, which is an established PLD regulator (Galas *et al*., 1997; Caumont *et al*., 1998, 2000; Yang and Mueckler, 1999; Vitale *et al*., 2002a, 2002b; Matsukawa *et al*., 2003; Liu *et al*., 2005; Béglé *et al*., 2009; Pelletán *et al*., 2015; Wang *et al*., 2023). This interaction happens after stimulation, suggesting that Ca^2+^ entry starts a cascade of events that leads to PA production, through these regulatory proteins. Therefore, PLD1 is present in positions where PA production could be needed, but only produces PA after activation.

Since PA is observed to be present post-fusion and PA is hypothesized as a lipid that stabilizes the fusion pore, we were curious if the amount of PA correlated to the rate of release as measured by the loss of VAMP2 from the fusion site. After binning fast fusion events from slow fusion events, more PA was observed at fast events (Figure 4I and S5). The role of PA in membrane fusion is likely coupled to the proteins it interacts with. In the case of PI (4,5)P_2_, more PI (4,5)P_2_ slowed fusion by recruiting endocytic proteins that restrict fusion pore expansion (Omar-Hmeadi *et al*., 2023). Similarly, PA could also act by interacting with proteins that assist with fusion and future studies could assess the proteins involved.

Overall, PLD1 has been established as necessary for stimulated exocytosis (Humeau *et al*., 2001; Vitale *et al*., 2001; Hughes *et al*., 2004; Jenkins and Frohman, 2005; Zeniou-Meyer *et al*., 2007). The results shown in this work support the hypothesis that PLD1 is involved due to its production of PA. PLD1 is present on vesicles, PA is formed after vesicles dock and PA accumulates at the fusion site post-fusion. In both cases, the formation of PA depends on PLD1 activity, which cannot be compensated by PLD2.

## METHODS

### Cell Culture

PC12-GR5 cells (gift from Dr. Wolf Almers) were cultured in flasks in Dulbecco’s modified eagle medium (DMEM, ThermoFisher, Waltham, MA, USA) supplemented with 5% fetal bovine serum (ThermoFisher, Waltham, MA, USA) and 5% equine serum (ThermoFisher, Waltham, MA, USA) and incubated at 37°C and 5% CO_2_. For imaging, PC12 cells were plated in 8 well plates (Cellvis, Mountain View, CA, USA) treated with poly-L-lysine (Sigma Aldrich, St. Louis, MO, USA). Cells were transfected with Lipofectamine 2000 (ThermoFisher, Waltham, MA, USA) and plasmids (25-100 ng/well) for fluorescently tagged proteins. The EGFP-PLD1 plasmid was a gift from Jeremy Baskin (Bumpus and Baskin, 2017). The PLD1 plasmid is a full-length version of PLD1 with an N-terminal EGFP tag. Specifically, GFP-PLD1 plasmid was made using the PLD1 ORF clone (NCBI accession # BC068976), amplifying it and inserting it into the GFP-C1 vector (Bumpus and Baskin, 2017). VAMP2-pHmScarlet was a gift from Pingyong Xu (Addgene plasmid # 166890). GFP-PASS was a gift from Guangwei Du (Addgene plasmid # 193970). Cells are tested for mycoplasma using the MycoFluor Mycoplasma Detection Kit (ThermoFisher, Waltham, MA, USA).

To stimulate fusion, a stimulation buffer containing 3 mM NaCl, 140 mM KCl, 1 mM MgCl_2_, 3 mM CaCl_2_, 10 mM D-glucose and 10 mM HEPES, pH 7.4 (ThermoFisher, Waltham, MA, USA) was added to a final concentration (KCl) of 60 mM. In inhibitory experiments, cells were incubated with 0.013% DMSO with or without 100 nM pan-PLD inhibitor 5-fluoro-2-indolyl des-chlorohalopemide (PLD1/2i, Sigma Aldrich, St. Louis, MO, USA) or 500 nM PLD1-specific inhibitor VU0155069 (PLD1i, Sigma Aldrich, St. Louis, MO, USA) for 30 minutes at 37°C and then imaged immediately. Knockdown experiments were transfected with 8 pmol siRNA (Scramble: AM4635, PLD1: 4390824, ThermoFisher, Waltham, MA, USA) and 0.4 µL Lipofectamine 2000 per well for 72 hours prior to imaging.

### Total Internal Reflection Fluorescence Microscopy

PC12 cells were imaged in Fluobrite DMEM (ThermoFisher, Waltham, MA, USA) using a TIRF microscope (Nikon Ti-U) equipped with 491 nm and 561 nm lasers, as described previously (Mahmood *et al*., 2023). A 60x 1.49 NA objective, a 2.5x magnifying lens, and an EMCCD (Andor iXon897, Abingdon, UK) were used in combination with a DualView (Optical Insights, Suwanee, GA, USA) to split the red and green fluorescence channels onto the camera via a 565LP dichroic with 525/50 and 605/75 emission filters (Chroma Technologies, Bellows Falls, VT, USA). Both color channels were taken simultaneously, and images were collected by MicroManager at 0.109 µm/pixel and 136 ms/frame (Edelstein *et al*., 2010). For tracking and inhibition studies, cells were imaged on a stage heater at 37°C. Otherwise temperature is noted in the figure captions for all data.

### Image Analysis

Image analysis was conducted using MATLAB (v. R2021b, Natick, MA, USA) To measure the colocalization of static images, red fluorescent objects in the Syx1a and granule marker images were located using a freely available spot finding routine (Crocker and Grier, 1996). All granules and Syx1a clusters found in the red channel were cropped in the corresponding green channel, making this and objective, object-based analysis, where the user does not choose what is colocalized and what is not. From the cropped green channel images, the, ΔF/S was calculated as follows: 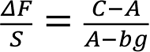, where C is the intensity of a 7 pixel circle in the green channel at a spot found in the red channel, A is a 1 pixel wide concentric ring with one pixel gap from the circle, and bg is the average intensity of the background surrounding the cell.

For measurements that take place in time, movies were corrected for photobleaching by a home-built code to correct to a constant cell intensity. Visiting, docking and fusing vesicles were found using a previously established algorithm (Mahmood *et al*., 2023). Here, a difference movie was calculated, max projected, bandpass filtered and a peak finding algorithm identify spots where a rapid increase of intensity occurs. Locations of intensity changes in the red channel were identified and cropped from the photobleach corrected movies (both green and red). From these cropped movies, intensity plots, calculated as 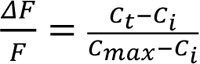, where C is the intensity of a 7 pixel circle as described in Figure 1 at frame t (C_t_) or the frame with the brightest intensity (C_max_), and C_i_ is the average initial intensity of a circle for 5 frames prior to onset of intensity increase. From the traces, data is manually binned into visiting, docking and fusing vesicles. If the trace is not clear, the cropped movie is viewed to determine the fate of the vesicle. Fusion is verified in the movie by an outward expansion of fluorescence post fusion. For average images of events, only events with at least 50 frames prior to event onset and 100 frames after were included. The bar graphs relating the ΔF/F for PLD1 or PASS intensity were measured at the time corresponding to the peak in intensity for each event for fusion and visiting vesicles. For the docking events, 80% to the plateau of the VAMP2 intensity was used instead of the peak intensity. To determine the relative rate of fusion upon treatment with siRNA, fusion events were identified as described above and counted in cells stimulated with 60 mM KCl, then divided by the average rate of fusion in control cells taken on the same day and from the same cell preparation. All movies were included, even if no fusion events were observed.

To determine the relative PASS intensity for the different stages of fusion the PASS intensity within a circular region divided by the average cell intensity was measured. This corrects for GFP-PASS protein expression levels. To obtain the relative PASS intensity for the visiting (orange), docking (pink) and fusion (blue), the PASS_initial_ intensity for visiting vesicles was subtracted. The formation or accumulation of PA (via PASS intensity) at fusion, docking and visiting sites is noted by an increase above 0 in the relative intensity and is compared to the PASS_initial_ intensity prior to vesicles visiting.

To determine the mobility of vesicles, tracking of SVs was done on VAMP2-pHmScarlet vesicles following a previously published analysis (Crocker and Grier, 1996). A position list of peaks was created for each frame, then a tracking algorithm determines tracks from those points. After tracking, the intensity (ΔF/S) of the PLD or PA in the vesicle’s position was measured in time using home-built code. The MATLAB code to locate and crop fusion, visiting and docking events is available on GitHub (https://github.com/michelleknowles/membrane-fusion) and previously published (Mahmood *et al*., 2023). JACoP in ImageJ was used to identify the percentage of Syx clusters or secretory vesicles that colocalized with PLD1 using the object-based methods (Bolte and Cordeliéres, 2006).

All statistical significance testing and plotting were conducted using appropriate tests in GraphPad Prism (v. 9.5.1, San Diego, CA, USA) according to published guidelines (Pollard *et al*., 2019). P-values were calculated by ANOVA followed by Tukey-Kramer post-hoc test when multiple comparisons were made (Figures 3A-C, 4G, 5D, and 6D), ANOVA followed by Donnett’s post-hoc test when multiple comparisons were made to one control (Figures 1D-E), or Student’s t-tests when only two things were compared (Figures 2G-I, 6C). Paired student’s t-tests were used when comparing the same data before and after an event like visiting, fusion, or docking (Figure 6D).

## Data availability

Data generated in this work will be made available from the corresponding author upon reasonable request.

## Code availability

Code for automatic detection of docking, fusion and visiting vesicles has been previously published (Mahmood *et al*., 2023) and is available on Github (https://github.com/michelleknowles/membrane-fusion). Any other code in this work will be made available from the corresponding author upon reasonable request.

## Acknowledgements

BLB acquired and analyzed data, contributed to writing and experimental design. MLH contributed to data analysis and data acquisition, MKK obtained funds, contributed to writing and data analysis. The work was funded by the National Science Foundation, Chemistry of Life Processes Grant # 1807455.

## Abbreviations

PLD: Phospholipase D

PA: Phosphatidic Acid

NPY: Neuropeptide Y

VAMP2: Vesicle Associated Membrane Protein 2

PABD: PA Binding Domain

