## Supplemental Materials for "Phospholipase D1 Produces Phosphatidic Acid at Sites of Secretory Vesicle Docking and Fusion"

*Supplementary Methods:* Calcium imaging was performed on a point-scanning confocal microscope (Olympus Fluoview 3000) with 488 and 640 laser excitation to observe a change in intensity post stimulation with high $K^+$. Cells were treated with Fluo 4 AM (ThermoFisher, Waltham, MA, USA) and far-red CellMask (ThermoFisher, Waltham, MA, USA) to test the influx of $Ca^{2+}$ upon addition of 3 mM NaCl, 140 mM KCl, 1 mM $MgCl_2$, 3 mM $CaCl_2$, 10 mM D-glucose and 10 mM HEPES, pH 7.4 (ThermoFisher, Waltham, MA, USA) to a final concentration (KCl) of 60 mM. CellMask and Fluo4-AM were used according to the manufacturer protocols.

To quantify slot blots, average intensities of each band were measured using Image J and a background intensity nearby was also measured. Values were corrected by normalizing to actin, then divided by the value of wild type cell from the same day. Specifically: $I_{Relative} = \frac{I_{O,PLD1} - I_{O,BG}}{I_{WT,PLD1} - I_{WT,BG}} * \frac{I_{WT,Actin} - I_{WT,BG}}{I_{O,Actin} - I_{O,BG}}$, where O is overexpressed, WT is wild type, PLD1 is the PLD1 band, Actin is the actin band and BG is a nearby background intensity surrounding the adjacent band.

For immunostaining, PC12 cells expressing VAMP2-pHluorin were fixed in 3.2% paraformaldehyde (Electron Microscopy Sciences, Hatfield, PA, USA) for 30 minutes and immediately permeabilized with 0.1% Triton-X 100 (Sigma Aldrich, St. Louis, MO, USA) for 10 minutes. Cells were then blocked in 0.5 mg/mL BSA (ThermoFisher, Waltham, MA, USA) for 4 hours at 4°C and treated with mouse anti-PLD1 (Santa Cruz Biotechnology, Santa Cruz, CA, USA) overnight. Cells were then treated with Alexa-594 conjugated rabbit anti-mouse (ThermoFisher, Waltham, MA, USA) for 4 hours. Transfection efficiency was 25-30%.

To determine the mobility of vesicles, tracking of SVs was done on VAMP2-pHmScarlet vesicles following a previously published analysis (Crocker and Grier, 1996). A position list of peaks was created for each frame, then a tracking algorithm determines tracks from those points. The diffusion coefficient (D) is calculated from individual tracks by calculating the mean squared displacement.

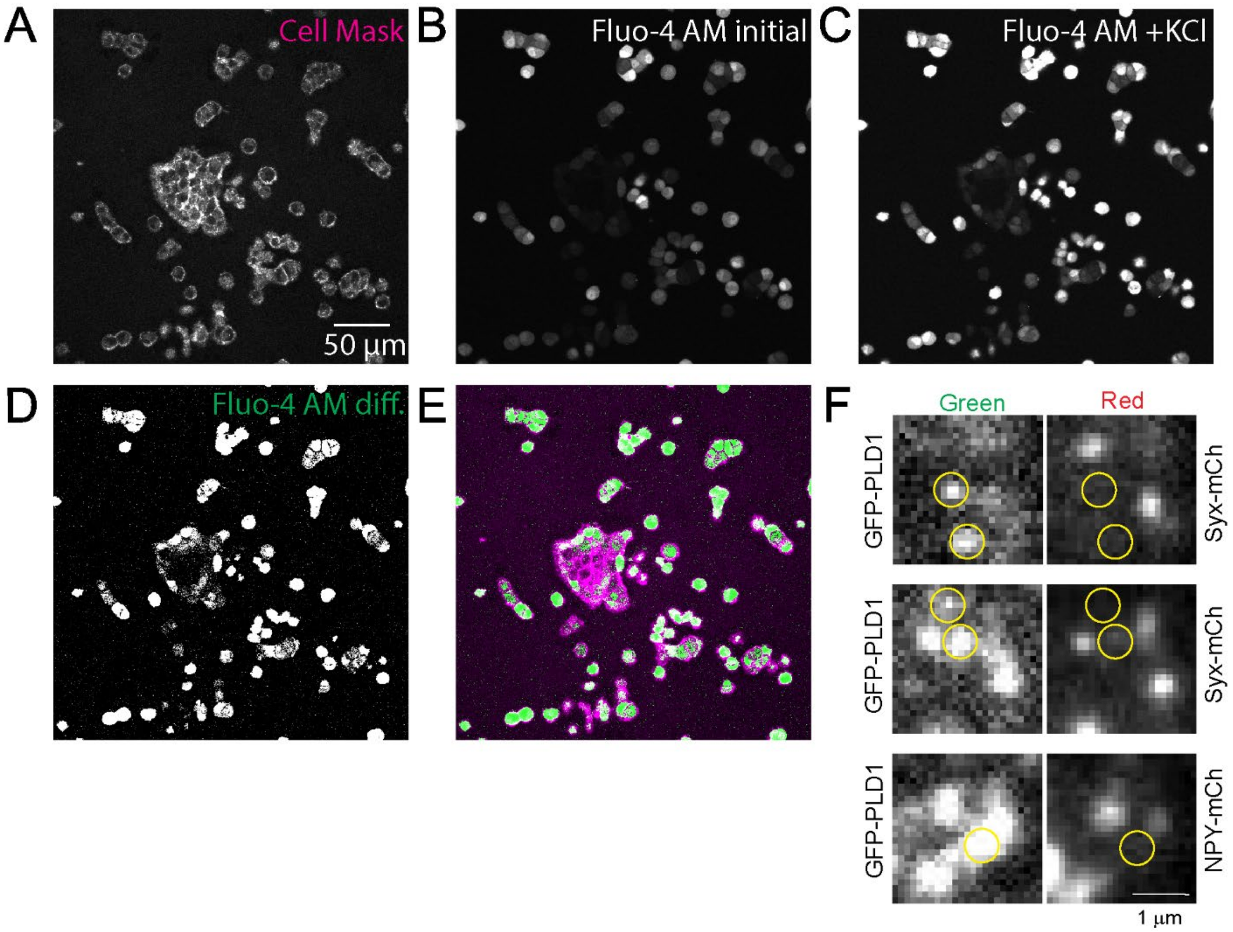


**Supplemental Figure 1: Controls for K+ stimulation and colocalization measurements.** A-E) Calcium influx occurs in KCl stimulated PC12 cells**.** PC12 cells were treated with Deep Red CellMask and the $Ca^{2+}$ dye Fluo-4 AM and imaged on a confocal microscope. Images were taken before and after the addition of stimulation buffer to a final KCl concentration of 60 mM. A) Average of 50 frames at 2.17 s/frame in CellMask channel. B-D) average of 5 frames prior to (B) or after (C) KCl stimulation and the difference between the images (D). B and C are contrasted identically. E) Composite image of the difference image shown in D (green) and the CellMask image shown in A (magenta). F) Bleed through of the green into the red channel is not observed. Locations where bright green spots were observed (left), do not always correspond to fluorescence in the red channel (right). The yellow circles indicate where GFP-PLD1 is bright but no red spot is visible in the secretory marker channel (Syx1a-mCherry or NPY-mCherry). The images shown here have been cropped from the images shown in Figure 1 (main paper). The cropped images have been rescaled.

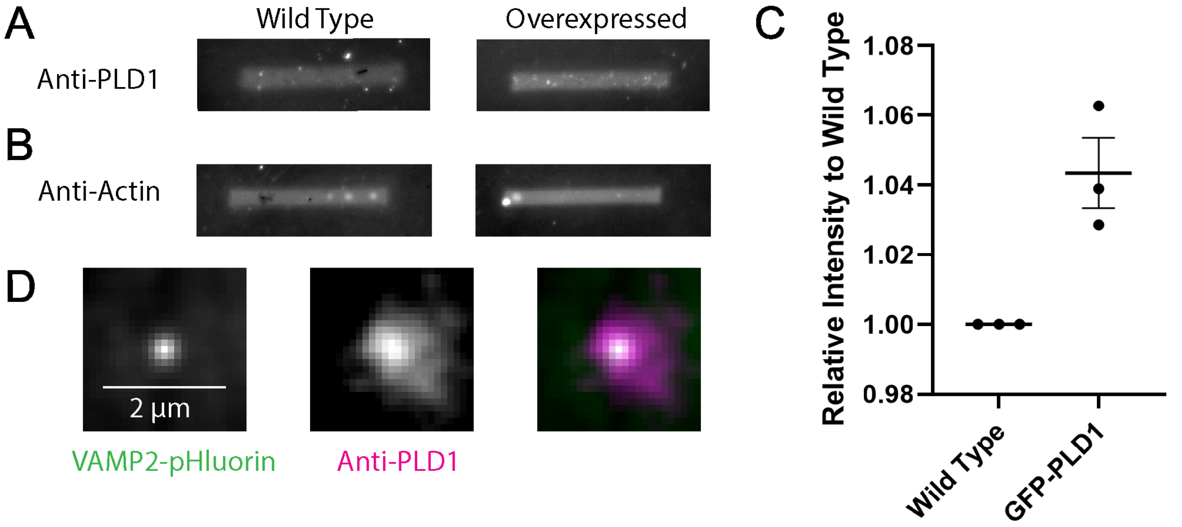


**Supplemental Figure 2: Blotting Overexpression of PLD1 and Staining for Localization**. A) Example Slot Blot of PLD1 and actin in wild type PC12 cells and PC12 cells transfected with GFP-PLD1. C) Normalized and corrected intensity of bands for PLD1. Values were corrected by normalizing to actin, then divided by the value of wild type cell from the same day. Lines are average, error bars are SEM (n = 3 days). C) Cells expressing VAMP2-pHluorin (left) were fixed, permeabilized and stained with anti-PLD1 and Alexa594-labeled anti-mouse (middle). On overlay of the two is also shown.

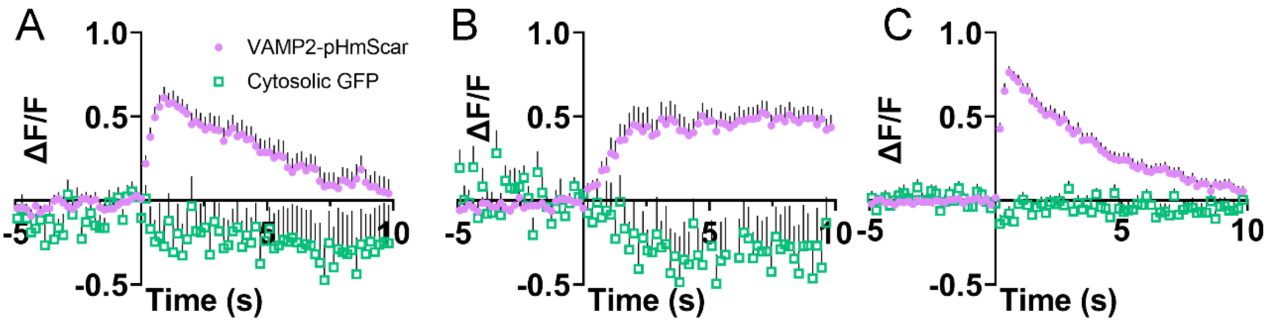


**Supplemental Figure 3: Cytosolic GFP does not increase during moving, docking or fusion events.** The normalized intensity of VAMP2-pHmScarlet (purple circles) and cytosolic GFP (green squares) during A) Visiting, B) Docking and C) Fusion events. Error is SEM.

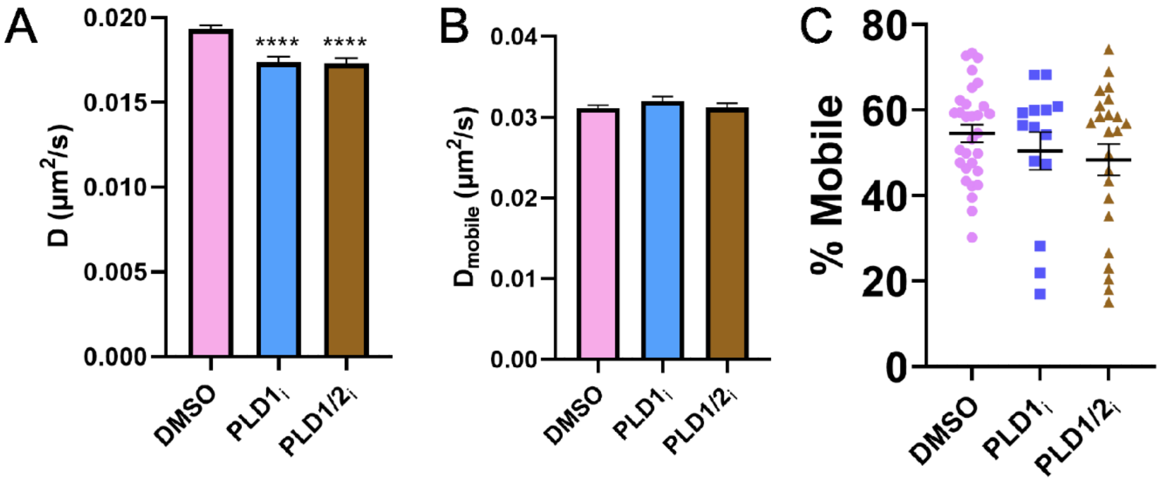


**Supplemental Figure 4: PLD1 inhibition reduces fraction of mobile vesicles.** A) Diffusion coefficients of single, tracked VAMP2 vesicles (n > 9500 tracks in at least 14 cells from at least 4 independent experiments) decreases slightly in the presence of inhibitors. B) Average diffusion coefficients of the mobile VAMP2 vesicles, where mobility is defined as D > 0.0055 $\mu m^2/s$. C) Percent of vesicles that are mobile in a given cell. Each point represents one cell from at least 4 independent experiments. Lines are averages, error bars are SEM. **** $p < 0.0001$ from an unpaired t-test compared to DMSO.

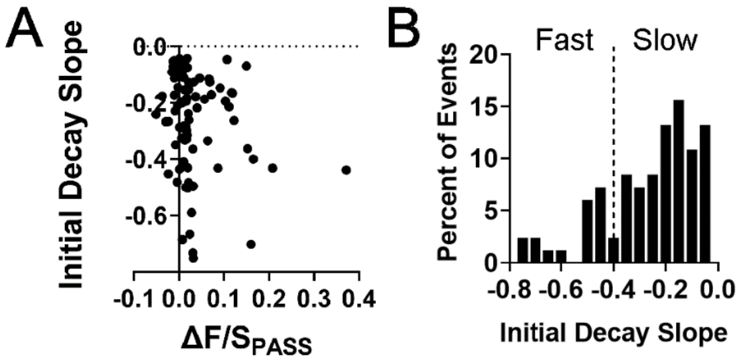


**Supplemental Figure 5: Correlation between the rate of decay and the amount of PA.** A) Initial decay slopes of VAMP2 and ΔF/S of GFP-PASS during fusion, every 10 events were averaged. B) Fast and slow fusion decays were sorted by the slope of the VAMP2 fusion decay as measured from the VAMP2 fluorescence peak to 1s later.

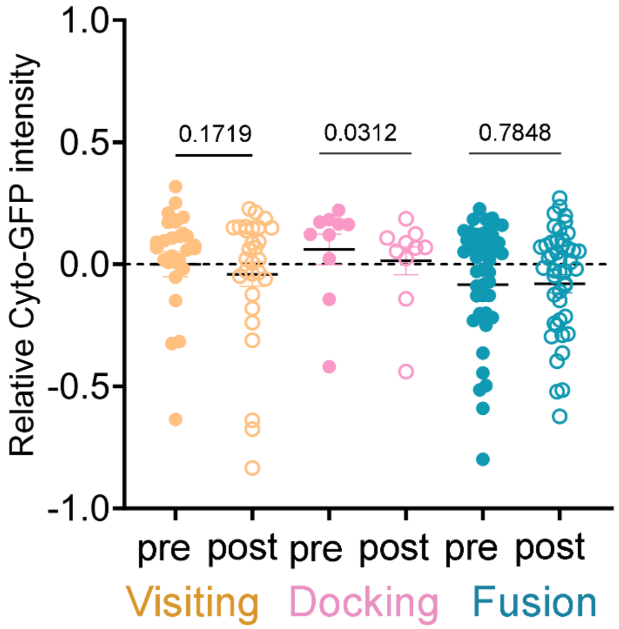


**Supplemental Figure 6: Relative change in cytoplasmic GFP intensity during visiting, docking, and fusion events.** Cytoplasmic GFP intensity over the different stages of membrane fusion: visiting (orange), docking (pink), and fusion (blue). The cytoplasmic GFP pre intensity (visiting) was subtracted from all to obtain a relative intensity. The "pre" intensity was measured at 0.5 to 0.1 s prior to the event. The post-docking intensity (pink, open circles) was measured after several seconds, when single events reached the plateau intensity in the VAMP2-pHmScarlet channel. The decrease in intensity during docking could be due to displacement of cytoplasmic GFP by secretory vesicles, which does not contain cytoplasmic GFP. The post-fusion intensity was measured at the peak of the VAMP2-pHmScarlet intensity for single fusion events. The p values represent paired t-tests for before and after values of the same event.
